# CD300a Blockade attenuates renal fibrosis by enhancing macrophage efferocytosis and neutrophil MMP9 production

**DOI:** 10.64898/2026.09.25.754396

**Authors:** Jinao Li, Chigusa Nakahashi-Oda, Hitoshi Koizumi, Fumie Abe, Kazuko Shibuya, Akira Shibuya

## Abstract

Renal fibrosis represents an irreversible endpoint of chronic kidney disease. Although macrophage efferocytosis regulates the response to acute renal injury, the mechanisms by which early efferocytosis confers long-term protection against chronic fibrosis remain elusive. Here, we show that phagocyte-specific deletion of the inhibitory receptor CD300a (*Cd300a*^fl/fl^*Lyz2-*Cre) mitigates obstructive renal injury and subsequent fibrosis. CD300a deficiency enhanced efferocytosis by tissue-resident macrophages during the early phase of injury, thereby suppressing the release of damage-associated molecular patterns (DAMPs) and TGF-β expression. This early attenuation of the inflammatory microenvironment altered the composition of recruited neutrophils, shifting them from profibrotic SiglecF+ toward non-profibrotic SiglecF− subsets. Furthermore, loss of CD300a cell-autonomously primed neutrophils for enhanced matrix metalloproteinase-9 (MMP9) production upon injury, whereas neutrophil depletion abolished the protective effect of CD300a deficiency. Importantly, administration of a neutralizing anti-CD300a antibody to wild-type mice recapitulated these protective effects, promoting the shift toward SiglecF− neutrophils and significantly attenuating renal fibrosis. Together, our findings identify CD300a as a regulator of early macrophage efferocytosis and neutrophil-derived MMP9 production, and establish pharmacological CD300a blockade as a potential therapeutic strategy for progressive renal fibrosis.

## Introduction

Chronic kidney disease (CKD) represents a major global health burden, with progressive renal fibrosis serving as the common pathological endpoint leading to irreversible organ failure (1–4). Following renal injury, massive stress and death of tubular epithelial and endothelial cells lead to the continuous release of damage-associated molecular patterns (DAMPs), such as high-mobility group box 1 (HMGB1) (5–7). These endogenous danger signals fundamentally alter the local tissue microenvironment, sustaining chronic inflammation and orchestrating cellular cascades that drive fibrogenesis (8). While early clearance of dead cells via efferocytosis is essential to limit DAMP leakage and reset the inflammatory microenvironment, the precise mechanisms that translate early efferocytosis into long-term attenuation of renal fibrosis remain poorly understood.

CD300a is an inhibitory immunoreceptor predominantly expressed on myeloid cells, including macrophages and neutrophils (9). Its cytoplasmic domain contains immunoreceptor tyrosine-based inhibitory motifs (ITIMs) that recruit SHP-1 and SHP-2 phosphatases to suppress activating signaling pathways (10). CD300a recognizes phosphatidylserine (PS) exposed on the outer membrane of dying cells and acts as a molecular brake on efferocytosis (11). We previously demonstrated that myeloid CD300a deficiency protects against acute organ injury in ischemia/reperfusion injury (IRI) models by enhancing macrophage-mediated efferocytosis and suppressing DAMP-induced inflammation (12). However, whether CD300a-mediated regulation of early efferocytosis and the post-injury microenvironment influences long-term progressive renal fibrosis—such as that induced by unilateral ureteral obstruction (UUO)—and how it modulates recruited immune cell responses remain unknown.

Recent studies highlight that tissue remodeling and fibrosis are dynamically regulated not only by resident macrophages but also by distinct heterogenous subsets of recruited neutrophils(12–15). In particular, a specific neutrophil subset marked by high expression of sialic acid–binding immunoglobulin-type lectin F (SiglecF) has been identified as a critical driver of a profibrotic microenvironment in injured tissues (16,17). However, whether CD300a influences the recruited neutrophil compartment or regulates the balance between profibrotic (SiglecF^+^) and non-profibrotic (SiglecF^−^) neutrophil subpopulations during renal fibrogenesis has not been established.

In this study, we investigated the pathophysiological role and therapeutic potential of CD300a in UUO-induced renal fibrosis. We show that phagocyte-specific CD300a deletion (*Cd300a*^fl/fl^*Lyz2*-Cre) enhances early efferocytosis by tissue-resident macrophages, suppressing initial DAMP release and TGF-*β* expression after UUO. This resolved early microenvironment, combined with cell-autonomous CD300a loss in neutrophils, drives a qualitative shift toward SiglecF^−^ neutrophils and markedly upregulates neutrophil-derived matrix metalloproteinase-9 (MMP9) to limit renal fibrosis. Furthermore, pharmacological blockade using a neutralizing anti-CD300a antibody successfully recapitulates this protection in wild-type mice. Collectively, our findings identify CD300a as a pivotal immunoregulatory checkpoint linking macrophage efferocytosis and neutrophil reprogramming to the resolution of progressive renal fibrosis.

## Methods

### Mice

C57BL/6J mice were purchased from Clea Japan (Tokyo, Japan). *Cd300a*^−/−^ (18), *Cd300a*^fl/fl^ (19), *Cd300a*^fl/fl^*Lyz2*-Cre mice(20) on the CS7BL/6 background were used. The mice used in the study were of the C57BL/6J background and were group-housed (3 to 5 mice per cage) in individually ventilated cages, following a 12-hour light-dark cycle. They were provided with unrestricted access to both food and water in a specific-pathogen-free environment. All animal experiments adhered to the guidelines set by the University of Tsukuba Animal Research Center’s animal ethics committee.

### Induction of UUO model

Male mice (8-12 weeks old, 23-29g) were used to induce UUO. Mice were anesthetized with an intraperitoneal injection of domitor (0.3 mg/kg), dormicum (4.0 mg/kg) and vetrophale (5.0 mg/kg). Kidneys were exposed through small abdomen incisions and ligate the proximal and distal segments of the right ureter with 5-0 silk thread under a binocular microscope, followed by a precise mid-segment incision (21). The abdomen and all layers of muscle and skin were closed with 6–0 nylon suture. After the surgical procedure, the mice were relocated to a clean cage positioned atop a heating pad maintained at a temperature of 42°C for 15min.

### Enzyme-linked immunosorbent assay (ELISA)

NGAL (Neutrophils Gelatinase-Associated Lipocalin) and HMGB-1 (High Mobility Group Box 1) levels in plasma were assessed using Mouse NGAL ELISA Kit (Proteintech) and HMGB1 ELISA Kit Exp (Shino-Test Corporation), following the respective manufacturer’s instructions.

### Biochemical examination

Serum BUN and creatinine were measured using FUJI DRI-CHEM SLIDE BUN-PIII (FUJIFILM) and CRE-PIII (FUJIFILM), following the respective manufacturer’s instructions.

### Cell isolation

To isolate renal cells, mice were deeply anesthetized and underwent transcardial perfusion with phosphate-buffered saline (PBS). Subsequently, the kidneys were harvested and placed in 5.0 mL of Dulbecco’s Modified Eagle Medium (DMEM) supplemented with 10% fetal bovine serum (FBS). The kidneys were then finely chopped using scissors and dissociated with a tumor dissociation kit (Miltenyi Biotec), following the manufacturer’s guidelines. After dissociation, erythrocytes were lysed using an ACK lysis buffer, and the resulting cell suspension was filtered through a 100 μm nylon mesh.

To isolate renal neutrophils, magnetic-activated cell sorting (MACS) (Miltenyi Biotec) was used by a negative selection strategy. Briefly, 10 μL of the renal cell suspension was taken for cell counting, followed by centrifugation at 300 g for 10 min at 4°C. The supernatant was removed, and cells were gently resuspended in 2% FBS in PBS buffer at 40 μL per 1x10⁷ cells. A neutrophil biotin-antibody cocktail (10 μL per 1x10⁷ cells) was added, and the suspension was gently mixed and incubated for 10 min at 2–8°C. Cells were then washed with 1–2 mL of 2% FBS in PBS buffer per 1x10⁷ cells and centrifuged. After removal of the supernatant, the pellet was resuspended in 80 μL of buffer per 1 x10⁷ cells, followed by the addition of anti-biotin microbeads (20 μL per 1x10⁷ cells) and incubation for 15 min at 2–8°C. Cells were washed again and resuspended in buffer at a concentration of up to 1x10⁸ cells per 500 μL. The cell suspension was applied to a MS column (Miltenyi Biotec) placed in a magnetic field, and the flow-through containing unlabeled neutrophils was collected. Columns were washed sequentially with buffer, ensuring that the column did not run dry. Collected cells were counted, centrifuged, resuspended at a final density of 2x10⁶ cells/mL, plated into 48-well plates (0.25 mL per well), and cultured under standard conditions (5% CO₂) with or without stimulation for 24h, flowed by flow cytometric analysis of SiglecF⁺ and SiglecF⁻ neutrophil populations or gene expression.

### Antibody and flow cytometry analysis

Cells were incubated with anti-mouse CD16/32 (2.4G2, BD Biosciences) in staining medium (2% FBS and 0.01% sodium azide in PBS) for 10 min to block the Fcγ receptors. Cells were then resuspended in the desired antibody and stained for 20 min on ice. All samples were analyzed by using an LSR Fortessa flow cytometer (BD Biosciences) followed by FlowJo software (TreeStar) and IDEAS6.2 (Merck), respectively. Fluorochrome-conjugated mAbs specific for CD45.2 (104), CD11b (M1/70), Ly6G (1A8), Ly6C (HK1.4), F4/80 (BM8) were purchased from BioLegend and SiglecF (E50-2440) was purchased from BD Biosciences. Anti-mouse CD300a mAb (EX42) was generated in our lab (22). Dead cells were excluded by using Zombie violet (423114, BioLegend) or propidium iodide (Sigma-Aldrich). Doublets were removed by excluding the population characterized by forward scatter and side scatter in flow cytometry throughout all experiments.

### Tissue preparation and histology

The kidneys were preserved using a 10% formalin-buffered solution, and 3 to 4 μm-thick slices were then embedded in paraffin. Following embedding, the paraffin-embedded tissue sections underwent staining with Masson’s trichrome or Sirius-red. Analysis was conducted utilizing a digital Keyence BZ-X700 light and fluorescent microscope. To assess fibrosis score, five fields were examined at 20 times magnification on each slide, and the final score was determined by calculating the average of these observations.

### Immunohistochemistry

In the context of multiplex immunohistochemistry (IHC), the staining process followed the Opal 7-Color Manual IHC Kit from PerkinElmer, adhering to the manufacturer’s instructions. Briefly, kidney sections on slides underwent a series of steps: baking at 65 °C for 1 hour, dewaxing with xylene, and dehydration with graded ethanol solutions. To block endogenous peroxidase activity, the samples were treated with 3% hydrogen peroxide for 15 minutes at room temperature (RT). Antigen retrieval was achieved using AR6 buffer (PerkinElmer) at 95-100 °C for 15 minutes, followed by incubation with Blocking One Histo (Nacalai Tesque) for 10 minutes at RT. Sections were incubated with polyclonal antibodies against kim-1 (1:200 dilution) (R&D, AF1817) and p-H3 (1:200 dilution) (Abcam, ab5176) in antibody diluent with background-reducing components (Dako) for 1 hour at RT or overnight at 4 °C. After three washes with Tris-buffered saline containing 0.05% Tween-20 (TBST), the slides were incubated with anti-mouse and rabbit Opal Polymer HRP (ARH1001EA; PerkinElmer) for 60 minutes at RT. Visualization using tyramide signal amplification (TSA) was carried out with Opal 570 (KIM-1). Subsequently, the slides were counterstained with the nuclear counterstain 4,6-diamidino-2-phenylindole (DAPI) (VectaShield, Vector Laboratories). The analysis of slides was performed using a Mantra 1.0.2 (PerkinElmer) fluorescence microscope, employing filters for Cy3, Texas Red and DAPI. This facilitated imaging for subsequent color unmixing based on their respective spectra. Unmixing of each OPAL fluorochrome and DAPI was done using mono fluorescence-stained slides, and multi-stained images underwent analysis with inForm Advanced Image Analysis software (inForm 2.4.1, PerkinElmer). For each sample, at least five fields at 20X magnification were analyzed, involving tissue segmentation, cell segmentation, and positive scoring.

### Efferocytosis assay

To assess the uptake of apoptotic cells by phagocytes, PSVue 643 (Molecular Targeting Technologies) was administered intravenously. At 0 hours, UUO surgery was performed, followed by the systemic injection of PSVue 643. After 1.5 hours, kidneys were harvested, and single-cell suspensions were prepared. Cells were subsequently analyzed by flow cytometry to evaluate PSVue+ dead cells and their engulfment by phagocytes.

### Neutrophil depletion

*InVivo*MAb anti-mouse Ly6G and *InVivo*MAb polyclonal rat IgG (control) were obtained from BioXCell. Mice were administered *InVivo*MAb anti-mouse Ly6G or *InVivo*MAb polyclonal rat IgG for three consecutive days with UUO surgery performed on the second day of antibody administration, designed as d 0. Kidneys were harvested for histology on day14.

### Quantitative RT-PCR

Total RNA was extracted from renal cells 3 day after UUO. RNA was extracted using ISOGEN (Nippon Gene). cDNA was synthesized using a High-Capacity cDNA Reverse Transcription Kit (Applied Biosystems). Quantitative RT-PCR (qRT-PCR) was performed using Power SYBER Green PCR Master Mix (Applied Biosystem) and an ABI 7500 sequence detector (Applied Biosystem). Normalization of quantitative real-time PCR was performed based on the gene encoding β-actin. The sequences of primers are shown, *β-actin* forward, 5’-ACTGTCGAGTCGCGTCCA 3’; *β-actin* reverse, 5’-GCAGCGATATCGTCATCCAT-3’; *Tgfb1* forward, 5’-TGACGTCACTGGAGTTGTACGG-3’; *Tgfb1* reverse, 5’-GGTTCATGTCATGGATGGTGC-3’; *Il6* forward, 5’-AGTTGCCTTCTTGGGACTGA-3’; *Il6* reverse, 5’-TCCACGATTTCCCAGAGAAC-3’; *Tnf* forward, 5’-AGTTGCCTTCTTGGGACTGA-3’; *Tnf* reverse, 5’-TCCACGATTTCCCAGAGAAC-3’; *Mmp9* forward, 5′-ACGACATAGACGGCATCCAGTATC-3′; *Mmp9* reverse, 5′-AGGTATAGTGGGACACATAGTGGG-3′; *Mmp8* forward *; Mmp8* reverse *;Prok2* forward, 5′-GCCCCGCTACTGCTACTTC-3′; *Prok2* reverse, 5′-CCGCACTGAGAGTCCTTGTC-3′; *Chil1* forward, 5′-GTACAAGCTGGTCTGCTACTTC-3′; *Chil1* reverse, 5′-ATGTGCTAAGCATGTTGTCGC-3′;.

### In vivo blocking of PS interaction with CD300a

D89E-MFG-E8 and the control protein (EPT-MFG-E8) (100 ug of each protein), which were generated as previously described (20), were administered intravenously 10 min before UUO.

### Preparation of the supernatant from kidney

The kidneys were harvested one day after UUO and finely minced using scissors. The tissue suspension was centrifuged, and the supernatant was collected and passed through a 70 μm nylon mesh. The filtrate was further aspirated using a 10mL syringe fitted with an 18-G needle, followed by filtration through a 0.2μm membrane filter. The filtered supernatant was then transferred to Amicon Ultra-4 centrifugal filter units with Ultracel-3K membranes (Millipore, Merck, Darmstadt, Germany) and centrifuged at 7,500 × g for 20 min (centrifugation conditions were adjusted as needed). Protein concentration was subsequently determined.

### Quantification and statistical analysis

Statistical significance of differences between groups was analyzed using GraphPad Prism 9 or 10 (GraphPad Software). The results are expressed as mean ± standard error of the mean (SEM). Statistical analyses were performed using one-way analysis of variance (ANOVA), followed by Šídák’s multiple-comparisons test for preselected pairwise comparisons. A *P*-value < 0.05 was considered statistically significant.

## Results

### CD300a deficiency alleviates tubular cell injury and fibrosis

We previously reported that macrophage efferocytosis at the very early stage of acute renal failure sequentially regulates chronic renal dysfunction (12). Indeed, following acute kidney injury, CD300a deficiency enhances macrophage efferocytosis within hours. To elucidate the downstream cellular mechanisms by which augmented efferocytosis actively suppresses collagen deposition during renal fibrosis, we generated phagocyte-specific CD300a-deficient mice (*Cd300a*^fl/fl^*Lyz2-* Cre mice) and subjected them to unilateral ureteral obstruction (UUO) (Fig. 1A). This well-established model induces tubular cell stress and death, recapitulating key features of human chronic obstructive nephropathy accompanied by progressive renal fibrosis. The levels of neutrophil gelatinase–associated lipocalin (NGAL), a marker released from damaged tubular epithelial cells (23), were markedly increased 24 h after UUO. Notably, NGAL concentrations were significantly lower in *Cd300a*^fl/fl^*Lyz2-*Cre mice than in control *Cd300a*^fl/fl^ mice (Fig. 1B). In contrast, blood urea nitrogen (BUN) and plasma creatinine levels were comparable between the two groups from 24 h to 14 days post-UUO (Supplementary Fig. 1A), likely because the functional contralateral kidney compensates for overall renal clearance. Nevertheless, at 14 days after UUO, expression of kidney injury molecule-1 (KIM-1) in the cortex of the obstructed kidney was markedly reduced in *Cd300a*^fl/fl^*Lyz2-*Cre mice compared with *Cd300a*^fl/fl^ mice (Fig. 1C and D). Collectively, these findings indicate that CD300a deficiency in phagocytes attenuates tubular cell injury, even though systemic renal function parameters remain unaffected due to contralateral compensation.

**Fig. 1.**
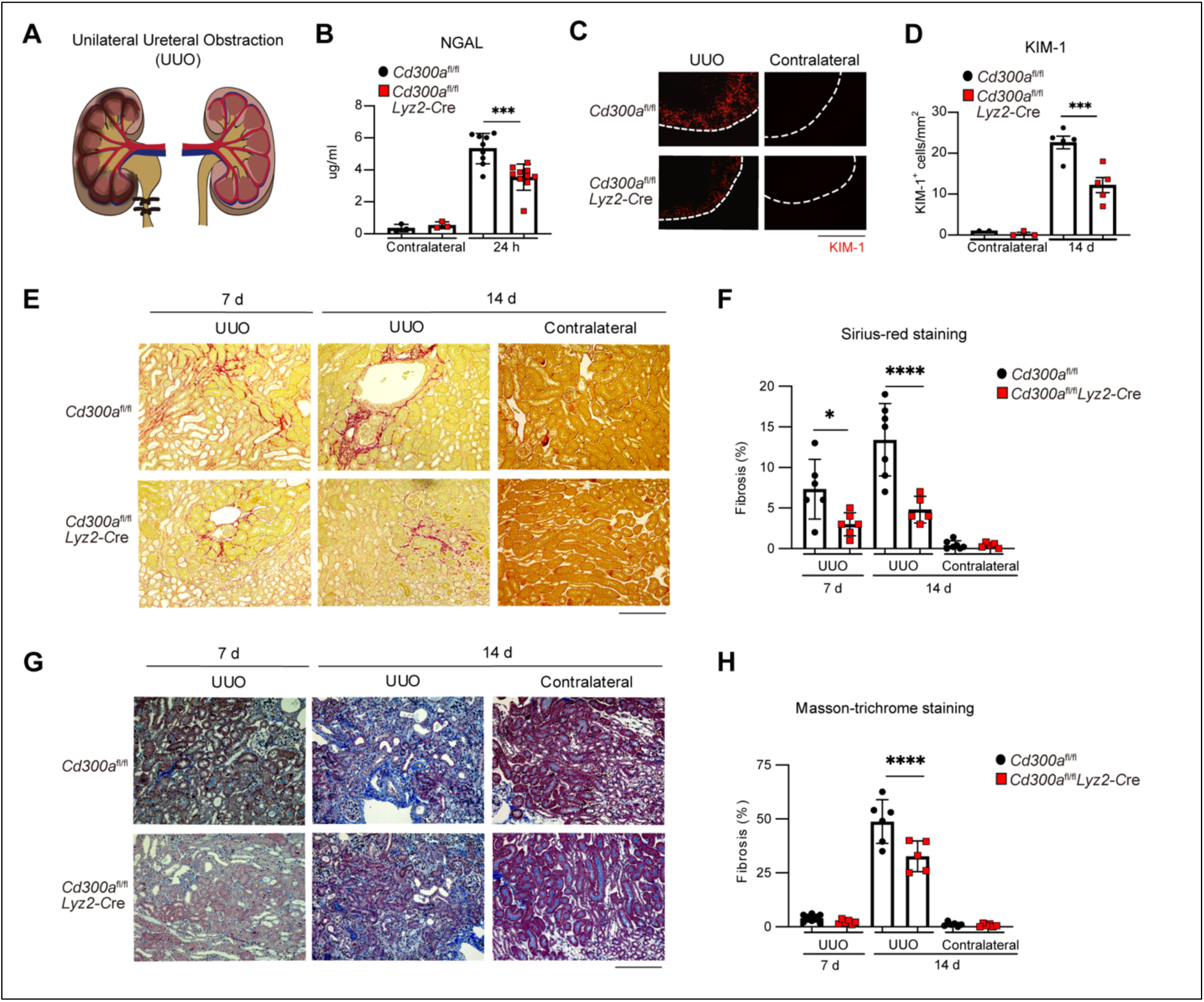
CD300a deficiency ameliorates tubular cell injury and reduces renal fibrosis. (A) Schematic diagram of Unilateral Ureteral Obstruction (UUO) model. (B) Plasma NGAL concentration in the contralateral and UUO side of kidney 24 h after UUO in *Cd300a*^fl/fl^ and *Cd300a*^fl/fl^*Lyz2-*Cre mice. (C and D) Immunofluorescence images of tubular cells stained with KIM-1 in *Cd300a*^fl/fl^ and *Cd300a*^fl/fl^*Lyz2-* Cre mice 7 and 14 d after UUO. (E–H) Representative Sirius-red (E) and Masson-trichrome (G) staining of kidneys on 7 and 14 d after UUO. Quantification of fibrosis area stained by Sirius-red (F) and Masson-trichrome (H). Scale bars, 100 µm. Data are presented as mean ± SEM. Statistics analyses performed using one-way ANOVA (B, D, F and H). *, *P* < 0.05, ***, *P* < 0.001, ****; *P* < 0.0001.

Although the renal function was not overtly impaired, the percentage of fibrotic area in the obstructed kidney progressively increased at 7 and 14 days post-UUO compared with the contralateral kidney, as evaluated by Sirius Red and Masson’s trichrome staining in five randomly selected fields of the cortico-medullary junction containing tubular cells (Supplementary Fig. 1B). Notably, renal fibrosis was significantly attenuated in *Cd300a*^fl/fl^ *Lyz2-*Cre mice compared with control *Cd300a*^fl/fl^ mice at both 7 and 14 days after UUO (Fig. 1E–H). Altogether, these data establish a crucial role for phagocytic CD300a in exacerbating tubular injury and renal fibrosis, prompting us to further investigate the underlying cellular and molecular mechanisms.

### Enhanced efferocytosis by CD300a-deficient macrophages suppresses renal inflammation and subsequent fibrosis

To clarify whether efferocytosis serves as a critical mechanism to suppress sequential fibrotic changes following tissue damage, we first confirmed the presence of dead cells in the kidney after UUO. Because dying cells lose the ability to maintain phosphatidylserine (PS) asymmetry, leading to PS exposure on the outer leaflet of the plasma membrane, we intravenously administered either the near-infrared fluorescent PS probe PSVue-643 or a PS-binding mutant protein of milk fat globule-EGF factor 8 (D89E-MFG-E8), which binds PS but lacks αvβ3integrin-binding capability, into UUO mice. Both approaches confirmed that PS-expressing dead cells were markedly induced by ureteral obstruction (Supplementary Fig. 2A).

To examine whether CD300a regulates efferocytosis post-UUO, we intravenously injected PSVue-643 into *Cd300a*^fl/fl^ and *Cd300a*^fl/fl^*Lyz2-*Cre mice immediately after UUO and evaluated efferocytosis in phagocyte populations via flow cytometry (Fig. 2A). *Cd300a*^fl/fl^*Lyz2-*Cre mice displayed a significantly higher proportion of PSVue-643-positive tissue-resident macrophages compared with control *Cd300a*^fl/fl^ mice at 1.5 h post-UUO. In contrast, neutrophils and inflammatory monocyte-derived macrophages exhibited minimal PSVue-643 fluorescence in both genotypes (Fig. 2B). These results indicate that phagocyte-specific deletion of CD300a selectively enhances efferocytosis by tissue-resident macrophages in the early phase following UUO.

**Fig. 2.**
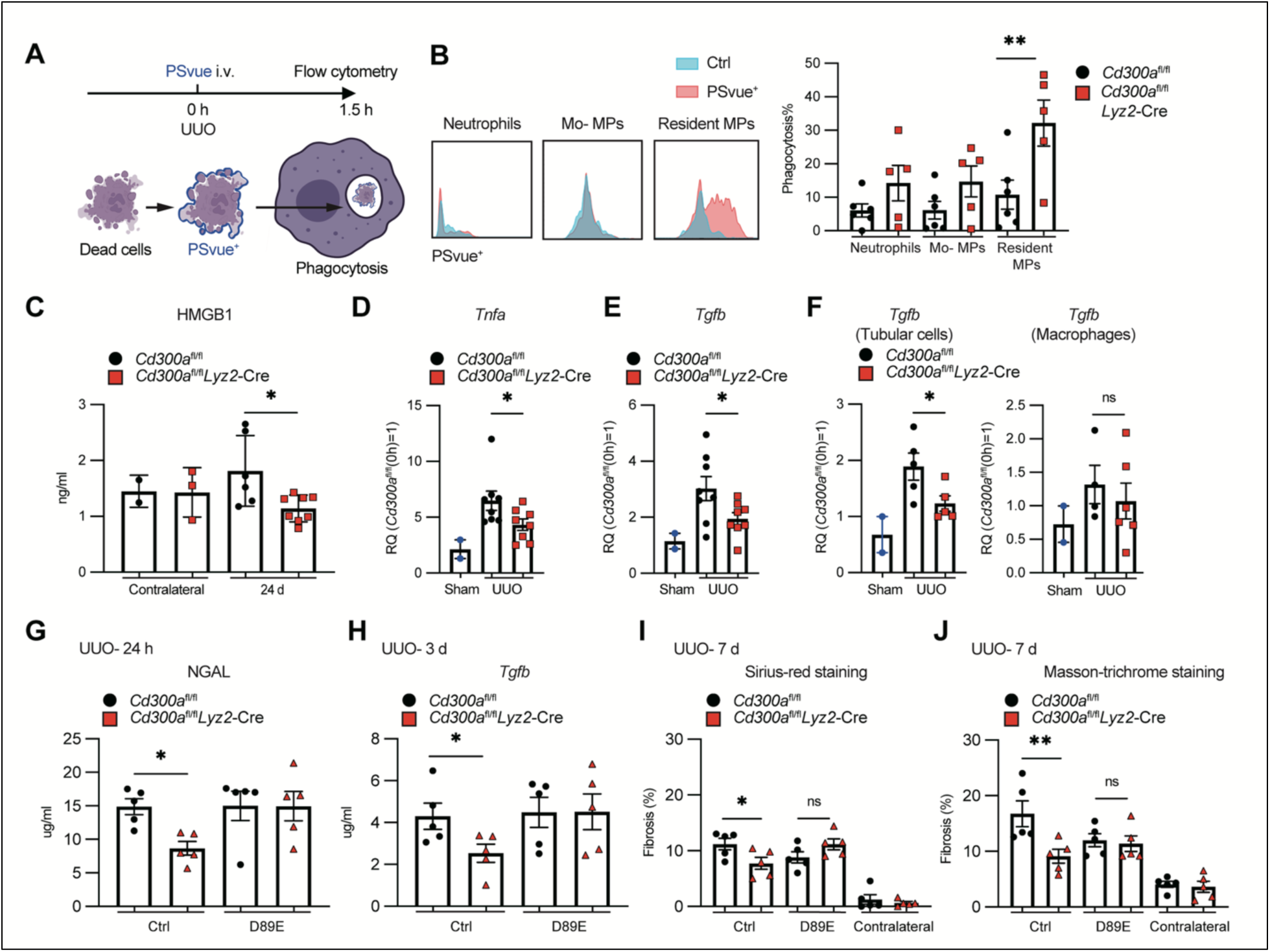
CD300a-deficiency enhanced efferocytosis by infiltrating resident macrophages. (A) Schematic diagram of efferocytosis analysis in the kidney after UUO. PSVue was intravenously injected just after UUO (0 h), and PSVue^+^ phagocytes were analyzed 1.5 h after UUO by flow cytometry. (B) Representative histograms of PSVue expression (left) and the percentages of PSVue^+^ cells in CD11b^+^ myeloid cells such as Ly6G^+^ neutrophils, CD11b^hi^ F4/80^int^ monocyte-derived macrophages (Mo-MPs) and CD11b^lo^ F4/80^int^ resident macrophages (resident MPs) in the kidney of *Cd300a*^fl/fl^ and *Cd300a*^fl/fl^*Lyz2-*Cre mice. (C) HMGB1 measured in the contralateral and UUO side of kidney 24 h after UUO in *Cd300a*^fl/fl^ and *Cd300a*^fl/fl^*Lyz2-*Cre mice. (D–F) Q-PCR analysis of *Tnfa* (D) and *Tgfb* expressions in the kidneys (E), isolated tubular cells and isolated macrophages (F) 3 d after UUO in *Cd300a*^fl/fl^ and *Cd300a*^fl/fl^*Lyz2-*Cre mice. (G) Plasma NGAL is measured by ELISA 24 h after UUO in EPT-MFG-E8 (Ctrl) or D89E-MFGE8 (D89E) administered *Cd300a*^fl/fl^ and *Cd300a*^fl/fl^*Lyz2-*Cre mice. (H) *Tgfb* expression at 3 d after UUO in Ctrl or D89E-administered *Cd300a*^fl/fl^ and *Cd300a*^fl/fl^*Lyz2-*Cre kidneys. (I and J) Quantification of Sirius-red (I) or Masson-trichrome (J) staining in kidneys from *Cd300a*^fl/fl^ and *Cd300a*^fl/fl^*Lyz2-*Cre mice administered with Ctrl or D89E at 7 d after UUO. Data are presented as mean ± SEM. Statistics analyses performed using one-way ANOVA (B–I). *, *P* < 0.05; **; *P* < 0.01.

UUO-induced death of endothelial and tubular epithelial cells promotes the release of damage-associated molecular patterns (DAMPs), thereby amplifying downstream inflammatory responses (12). Among these, HMGB1 is a prototypical DAMP that exacerbates kidney injury via TLR4 signaling (28). Consistent with enhanced clearance of dead cells, plasma levels of HMGB1 and renal expression of the proinflammatory cytokine TNF-α were significantly reduced in *Cd300a*^fl/fl^*Lyz2-*Cre mice compared with *Cd300a*^fl/fl^ mice at 24 h and 3 days post-UUO, respectively (Fig. 2C and D). In line with these findings, *Cd300a*^fl/fl^*Lyz2-*Cre mice showed diminished expression of transforming growth factor-β (*Tgfb*) in the obstructed kidney, suggesting that CD300a deficiency dampens inflammation during early tissue injury (Fig. 2E). Notably, *Tgfb* expression was significantly decreased in tubular epithelial cells isolated from *Cd300a*^fl/fl^*Lyz2-*Cre mice, whereas no significant difference was observed in kidney macrophages between genotypes (Fig. 2F). In contrast, *Il6, Il10* and *Il1b* expression levels were comparable between *Cd300a*^fl/fl^ and *Cd300a*^fl/fl^*Lyz2-*Cre mice’s kidneys (Supplementary Fig. 2B).

Wild-type MFG-E8 mediates efferocytosis by bridging PS on dead cells to α integrin on phagocytes. Conversely, mutant MFG-E8, D89E-MFG-E8 competes for PS binding without triggering integrin engagement, thereby dominant-negatively inhibiting PS-mediated efferocytosis (29). To confirm that the protective phenotype depends on efferocytosis, D89E-MFG-E8 or a control mutant protein (EPT-MFG-E8) was intravenously injected into *Cd300a*^fl/fl^ and *Cd300a*^fl/fl^*Lyz2-*Cre mice 10 min prior to UUO. Following control EPT-MFG-E8 administration, *Cd300a*^fl/fl^*Lyz2-*Cre mice retained significantly lower plasma NGAL levels and reduced renal *Tgfb* expression compared with *Cd300a*^fl/fl^ mice. Crucially, administration of D89E-MFG-E8 completely abolished these protective differences, yielding comparable NGAL and *Tgfb* levels between the two genotypes (Fig. 2G and H). In contrast, systemic parameters (BUN and plasma creatinine) remained comparable across all four groups from 24 h to 7 days post-UUO (Supplementary Fig. 2C). In agreement with the NGAL and *Tgfb* profiles, the attenuation of renal fibrosis observed in *Cd300a*^fl/fl^*Lyz2-*Cre mice was preserved upon EPT-MFG-E8 treatment but fully negated by D89E-MFG-E8 administration (Fig. 2I and J, Supplementary Fig. 2D and E). Collectively, these results demonstrate that CD300a deficiency in phagocytes enhances early efferocytosis by tissue-resident macrophages, which suppresses DAMP release and subsequent inflammatory and fibrotic cascades following obstructive renal injury.

### CD300a deficiency alters neutrophil subset composition and cell-autonomously upregulates MMP9 to attenuate post-obstructive renal fibrosis

After kidney injury, the innate immune response is rapidly activated, prompting the recruitment of circulating inflammatory immune cells into the damaged tissue (30). Although CD300a deficiency in phagocytes enhances early efferocytosis by tissue-resident macrophages and acutely suppresses initial inflammation, we reasoned that early clearance alone might not fully account for the long-term attenuation of renal fibrosis. Therefore, we next investigated the role of recruited inflammatory cells. The overall proportion and absolute numbers of recruited neutrophils and monocyte-derived macrophages were comparable between *Cd300a*^fl/fl^ and *Cd300a*^fl/fl^*Lyz2-*Cre mice following UUO (Supplementary Fig. 3A and B). Intriguingly, however, phenotypic analysis revealed a marked qualitative shift in neutrophil subpopulations. *Cd300a*^fl/fl^*Lyz2-*Cre mice displayed a significantly lower proportion of SiglecF⁺ neutrophils and a corresponding higher proportion of SiglecF⁻ neutrophils at 7 days post-UUO compared with *Cd300a*^fl/fl^ mice (Fig. 3A). To test whether this population shift was driven by the altered microenvironment of the injured kidney, bone marrow–derived neutrophils (BM neutrophils) from wild-type (WT) mice were stimulated in vitro with injured kidney supernatants obtained from either *Cd300a*^fl/fl^ and *Cd300a*^fl/fl^*Lyz2-*Cre UUO mice. WT BM neutrophils cultured with supernatants from *Cd300a*^fl/fl^*Lyz2-*Cre mice yielded a significantly higher proportion of SiglecF⁻ neutrophils than those cultured with supernatants from *Cd300a*^fl/fl^ mice (Fig. 3B). These results indicate that phagocyte-specific CD300a deficiency modifies the post-injury renal microenvironment to favor the induction or survival of the SiglecF⁻ neutrophil subset.

**Fig. 3.**
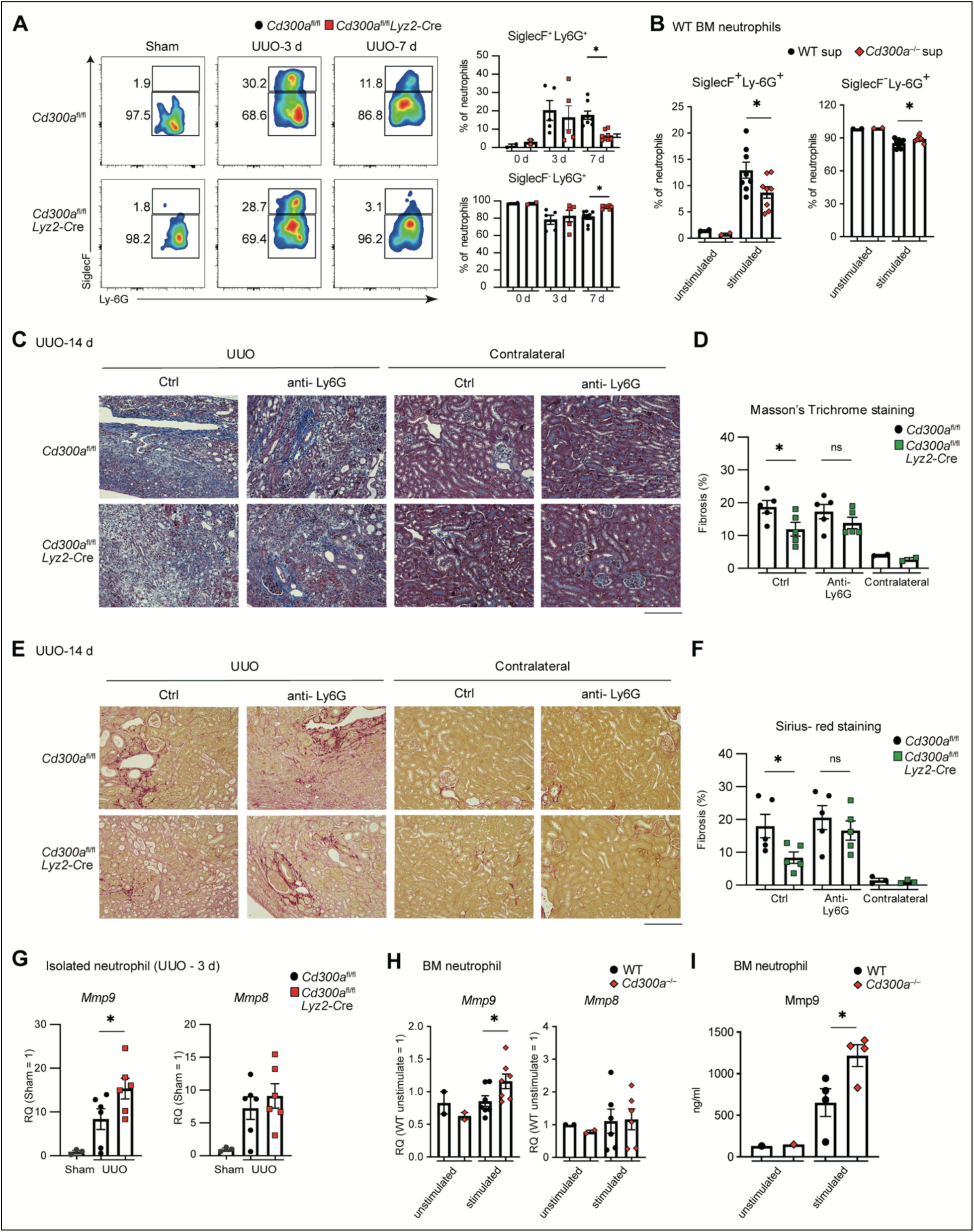
CD300a regulates the balance of SiglecF^+^ and SiglecF^-^ neutrophils in renal fibrosis. (A) Flow cytometric analysis of SiglecF⁺ and SiglecF⁻ neutrophils (left) and quantification of the percentages of SiglecF⁺ and SiglecF⁻ neutrophils in Sham (0 d) and post UUO day 3 (3 d) and day 7 (7 d) groups. (B) Bone marrow-derived (BM) neutrophils were isolated from wild-type (WT) mice and stimulated with kidney supernatants from WT or *Cd300a*^-/-^ mice after UUO for 24 h. Quantification of the percentages of SiglecF⁺ and SiglecF⁻ neutrophils with or without stimulation. (C to F) Representative Masson-trichrome (C) and Sirius-red staining (E) of ctrl Ab or neutrophil depleting, anti-Ly6G Ab (Anti-Ly6G), administered kidneys on 14 d after UUO. Quantification data is shown on the right representatively (D and F). (G) Gene expression levels of *Mmp9* and *Mmp8* in isolated neutrophils from *Cd300a*^fl/fl^ or *Cd300a*^fl/fl^ *Lyz2-*Cre mice at day 3 after UUO. (H and I) Bone marrow-derived neutrophils from WT and *Cd300a*^-/-^ mice were stimulated with kidney supernatants from WT mice after UUO for 24 h. Gene expression levels of *Mmp9* and *Mmp8* (H) by Q-PCR and Mmp9 concentration by ELISA(I). Data are presented as mean ± SEM. Statistics analyses performed using one-way ANOVA (A, B and D–I). *, *P* < 0.05.

A previous study demonstrated that SiglecF-expressing neutrophils play an essential role in orchestrating a profibrotic microenvironment during renal fibrosis (31). To assess whether neutrophils actively contribute to the CD300a-mediated protection against UUO-induced fibrosis, we depleted neutrophils using an anti-Ly6G monoclonal antibody (mAb) (Supplementary Fig. 3C). *Cd300a*^fl/fl^ and *Cd300a*^fl/fl^*Lyz2-*Cre mice were intravenously administered either anti-Ly6G mAb or control antibody on days −1, 0, and 1 relative to UUO surgery on day 0. In control antibody-treated mice, renal fibrosis following UUO was significantly reduced in *Cd300a*^fl/fl^*Lyz2-*Cre mice compared with *Cd300a*^fl/fl^ mice. Strikingly, neutrophil depletion blunted the difference in fibrosis between the two genotypes (Fig. 3C–F). These results demonstrate that the attenuated renal fibrosis in CD300a-deficient mice partially depends on neutrophils, indicating that altered neutrophil subset composition plays a central role in suppressing the profibrotic microenvironment.

To determine how neutrophils contribute to this protective effect, we examined whether neutrophils exhibit altered expression of fibrosis-associated genes following UUO. The expression of *Mmp9*, a matrix metalloproteinase known to regulate extracellular matrix remodeling and to be abundantly produced by activated neutrophils during tissue injury (32), was significantly elevated in neutrophils isolated from *Cd300a*^fl/fl^*Lyz2-*Cre mice compared with those from *Cd300a*^fl/fl^ mice at 3 days post-UUO. Although *Mmp8* expression showed an increasing trend, the difference did not reach statistical significance (Fig. 3G). Furthermore, the expression levels of other inflammatory and fibrotic mediators, including *Tnfa*, *Tgfb*, *Chil1*, and *Prok2*, were comparable between the two groups (Supplementary Fig. 3D).

To delineate whether this enhanced MMP9 expression reflects a cell-autonomous effect of CD300a loss in neutrophils, BM neutrophils from WT or CD300a-deficient (*Cd300a*^-/-^) mice were stimulated with the same injured kidney supernatants derived from WT UUO mice. Notably, *Cd300a*^-/-^ BM neutrophils exhibited significantly higher *Mmp9* mRNA expression and MMP9 protein production than WT BM neutrophils upon identical supernatant stimulation (Fig. 3H and I), whereas expression levels of *Mmp8*, *Tnfa*, *Tgfb*, *Chil1*, and *Prok2* remained unchanged (Supplementary Fig. 3E). Furthermore, CD300a deficiency in BM neutrophils did not alter the proportion of SiglecF⁺ or SiglecF⁻ subsets under identical supernatant stimulation conditions (Supplementary Fig. 3F). Collectively, these findings demonstrate that while the post-injury microenvironment in *Cd300a*^-/-^ mice dictates the shift toward the non-profibrotic SiglecF− neutrophil subset, CD300a intrinsically suppresses MMP9 production in neutrophils in a cell-autonomous manner. Thus, loss of CD300a mitigates renal fibrosis through the synergistic combination of a favorable renal microenvironment and cell-intrinsic upregulation of MMP9 in neutrophils.

### The blockade of CD300a ameliorates post-obstructive renal fibrosis

To evaluate the therapeutic potential of targeting CD300a in preventing renal fibrosis, WT mice were intravenously administered a neutralizing anti-CD300a mAb or an isotype control antibody 2 h prior to UUO surgery (Fig. 4A). Consistent with our findings in *Cd300a*^fl/fl^*Lyz2-*Cre mice (Fig. 1B and 3A), treatment with the anti-CD300a mAb markedly reduced plasma NGAL levels, decreased the proportion of SiglecF⁺ neutrophils, and concomitantly increased the non-profibrotic SiglecF⁻ neutrophil population (Fig. 4B and C). Moreover, histopathological analysis by Sirius Red and Masson’s trichrome staining demonstrated that renal fibrosis at 7 and 14 days post-UUO was significantly attenuated in anti-CD300a mAb-treated mice compared with control antibody-treated mice (Fig. 4D–G). In contrast, anti-CD300a mAb administration did not significantly affect systemic renal function markers (BUN or plasma creatinine) or alter the proportion of monocyte-derived macrophages and tissue-resident macrophages in the kidney at 3 or 7 days after UUO (Supplementary Fig. 4A–C). These data indicate that pharmacological blockade of CD300a successfully recapitulates the protective phenotype of genetic CD300a deficiency, highlighting anti-CD300a neutralizing antibodies as a promising therapeutic strategy to prevent renal fibrosis.

**Fig. 4.**
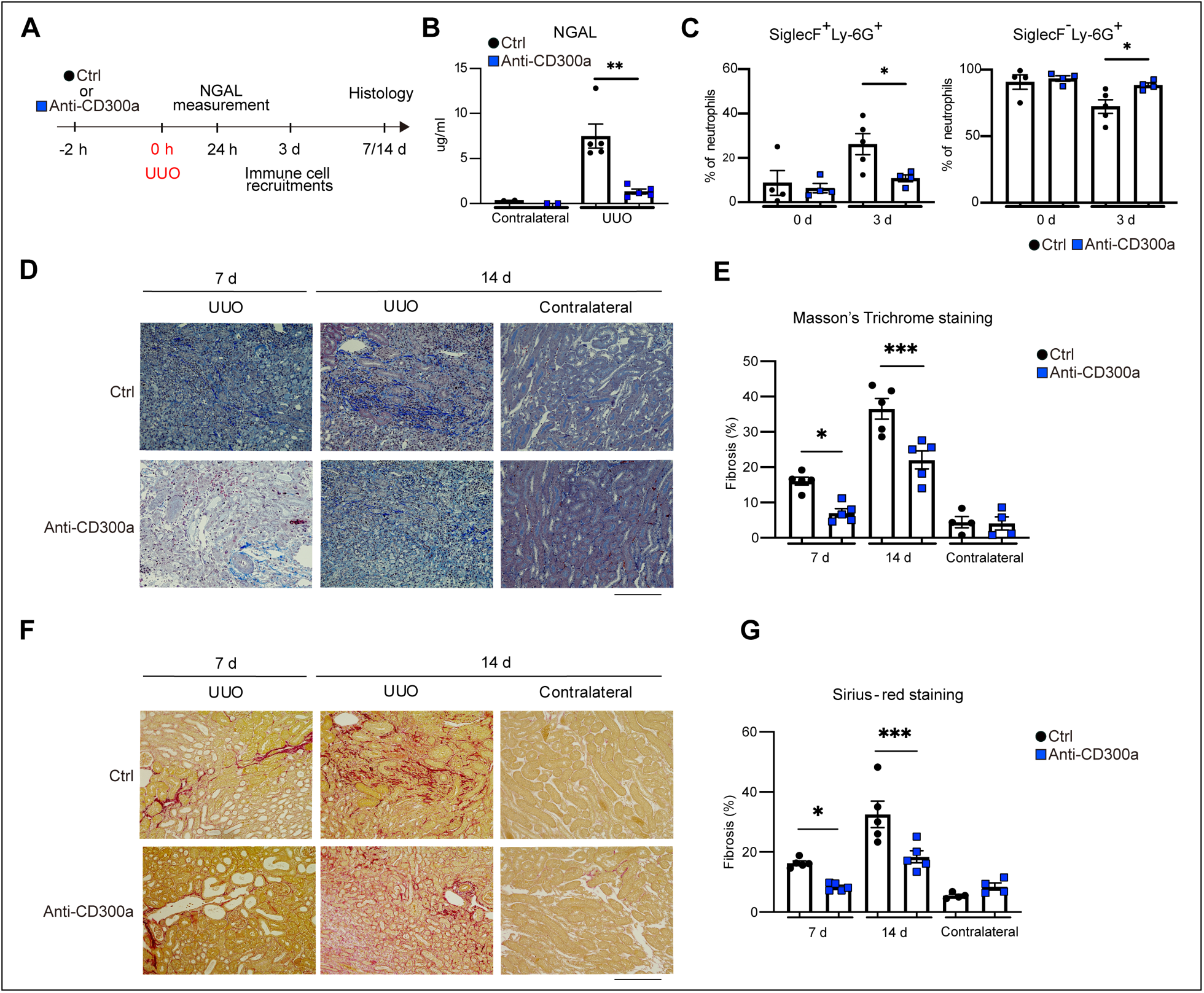
An anti-CD300a neutralizing mAb ameliorates tubular cell injury and renal fibrosis. (A) WT mice were treated with control antibody (Ctrl) or anti-CD300a antibody (Anti-CD300a) 2 h before UUO. Plasma NGAL levels were measured 24 h after surgery, and immune cell recruitment was analyzed at day 3. Kidneys were harvested on days 7 and 14 for assessment of renal fibrosis. (B) NGAL were measured on 24 h after UUO in Ctrl or Anti-CD300a treated mice. (C) Quantitative data of the percentage of SiglecF^+^ and SiglecF^-^ neutrophils in Ctrl or Anti-CD300a treated mice on 3 and 7 days after UUO. (D to G) Representative Masson-trichrome (D) and Sirius-red staining (F) of kidneys after Ctrl or Anti-CD300a treatment on day7 and day14 after UUO. Quantification data is shown on the right representatively (E and G). Scale bars, 100 µm. Data are presented as mean ± SEM. Statistics analyses performed using one-way ANOVA (B, C and E–G). *, *P* < 0.05; **, *P* < 0.01, ***, *P* < 0.001.

## Discussion

In the present study, we identify the inhibitory immunoreceptor CD300a as a pivotal immunoregulatory checkpoint linking early dead cell clearance to the resolution of progressive renal fibrosis. Using a phagocyte-specific CD300a-deficient mouse model (*Cd300a*^fl/fl^*Lyz2*-Cre) and a therapeutic neutralizing anti-CD300a antibody, we demonstrate that loss of CD300a orchestrates a dual protective mechanism against UUO-induced injury. Specifically, CD300a deficiency in tissue-resident macrophages enhances early efferocytosis, limiting DAMP release and suppressing profibrotic signaling in parenchymal cells. This reset microenvironment, operating in concert with cell-autonomous neutrophil reprogramming, drives a qualitative shift from profibrotic SiglecF^+^ toward non-profibrotic SiglecF^−^ neutrophils and selectively upregulates neutrophil-derived MMP9. Crucially, antibody-mediated CD300a blockade in wild-type mice recapitulates these protection mechanisms, providing compelling proof-of-concept for CD300a as a therapeutic target in chronic kidney disease.

Early tubular epithelial cell (TEC) injury and secondary necrosis are key initiating events in post-obstructive renal fibrogenesis (24,25). We observed that phagocyte CD300a deficiency markedly attenuated acute tubular stress, as evidenced by reduced plasma NGAL levels and decreased HMGB1 release shortly after UUO. Notably, *Tgfb* expression was selectively suppressed in TECs rather than in kidney macrophages. These findings support a model in which CD300a-deficient macrophages rapidly clear dying cells via enhanced efferocytosis, preventing secondary necrosis and DAMP leakage, which indirectly dampens stress-induced TGF-*β* production in neighboring parenchymal cells (26,27). Thus, CD300a acts as a molecular gatekeeper that regulates the early acute microenvironment to prevent chronic fibrotic priming.

A central finding of our work is the functional and phenotypic modulation of recruited neutrophils. Although overall neutrophil accumulation was comparable between genotypes, neutrophil depletion completely abrogated the anti-fibrotic protection conferred by CD300a deficiency, establishing neutrophils as indispensable mediators. Phenotypically, CD300a deficiency markedly reduced the profibrotic SiglecF^+^ neutrophil subset while increasing the SiglecF^−^ population (12). Functionally, *Mmp9* was selectively upregulated in kidney-isolated neutrophils from CD300a-deficient mice. The robust *Mmp9* induction observed *in vivo* compared with *in vitro* supernatant-stimulated BM neutrophils suggests that sustained exposure to the resolved renal microenvironment, combined with cell-intrinsic CD300a deficiency, synergistically enhances MMP9 production. Given that MMP9 facilitates extracellular matrix turnover and tissue remodeling during the resolution phase (28), this neutrophil-derived MMP9 axis actively accelerates matrix resolution rather than promoting tissue destruction.

Several limitations of the present study warrant consideration. First, while we established that CD300a deficiency cell-autonomously enhances neutrophil *Mmp9* expression, the precise intracellular signaling pathways downstream of CD300a—particularly how ITIM-mediated SHP- 1/2 recruitment intersects with transcriptional regulators of *Mmp9* in neutrophils—remain to be fully delineated. Second, although neutrophil depletion demonstrated an indispensable role for neutrophils in CD300a-mediated protection, definitive causal links between neutrophil-derived MMP9 upregulation and matrix degradation require further validation, such as *in vivo* MMP9 pharmacological inhibition or cell-specific gene deletion experiments. Finally, whether MMP9 upregulation is restricted to the SiglecF^−^ subset or universally induced across neutrophil subpopulations warrants detailed sorting and single-cell profiling. Addressing these mechanistic questions will further refine our understanding of CD300a as a therapeutic target in tissue remodeling.

From a translational perspective, demonstrating the therapeutic efficacy of an anti-CD300a neutralizing mAb represents a critical milestone. Systemic administration of the neutralizing mAb prior to obstructive injury successfully suppressed DAMP-induced inflammation, altered neutrophil subset dynamics (detectable by day 3 post-UUO), and attenuated renal collagen deposition without compromising overall renal function parameters. This rapid, disease-modifying effect highlights the feasibility of antibody-based intervention. By restoring endogenous efferocytosis and promoting a pro-resolving immune cell phenotype, CD300a targeting offers a unique immunotherapeutic approach that harnesses the innate repair machinery to halt fibrotic progression. In conclusion, our findings demonstrate that CD300a restrains tissue-resident macrophage efferocytosis and cell-intrinsically suppresses neutrophil MMP9 expression. Blockade of CD300a shifts the post-injury microenvironment toward tissue resolution, identifying CD300a as a highly attractive immunotherapeutic target to combat progressive chronic kidney disease.

## Acknowledgments

The authors thank M. Kaneko for administrative support.

## Author contributions

L. J. performed most experiments. H. K. and F. A. contributed to animal studies. L. J. and C. N-O. analyzed data. L. J., C. N-O., and A.S. wrote the manuscript. C. N-O., K. S. and A. S. supervised this project.

## Funding

This work was supported by the JSPS KAKENHI (21H04836 to AS, 24K02258 to KS, and 25K02740 and 26K23404 to C. N-O.) and by the Japan Agency for Medical Research and Development (AMED) under the Practical Research Project for Renal Diseases (Grant Number: 26ek0310026h0002 to AS).

## Conflict of interest

FA is an employee of TNAX Biopharma Corporation. KS and AS own a stake in TNAX Biopharma Corporation.

## Data availability

Source data will be provided upon reasonable request to the corresponding author.

