## Supplemental Figures for "CD300a Blockade attenuates renal fibrosis by enhancing macrophage efferocytosis and neutrophil MMP9 production"

Conflict of interest: FA is an employee of TNAX Biopharma Corporation. KS and AS own a stake in TNAX Biopharma Corporation.

Supplemental figures 1 – 4

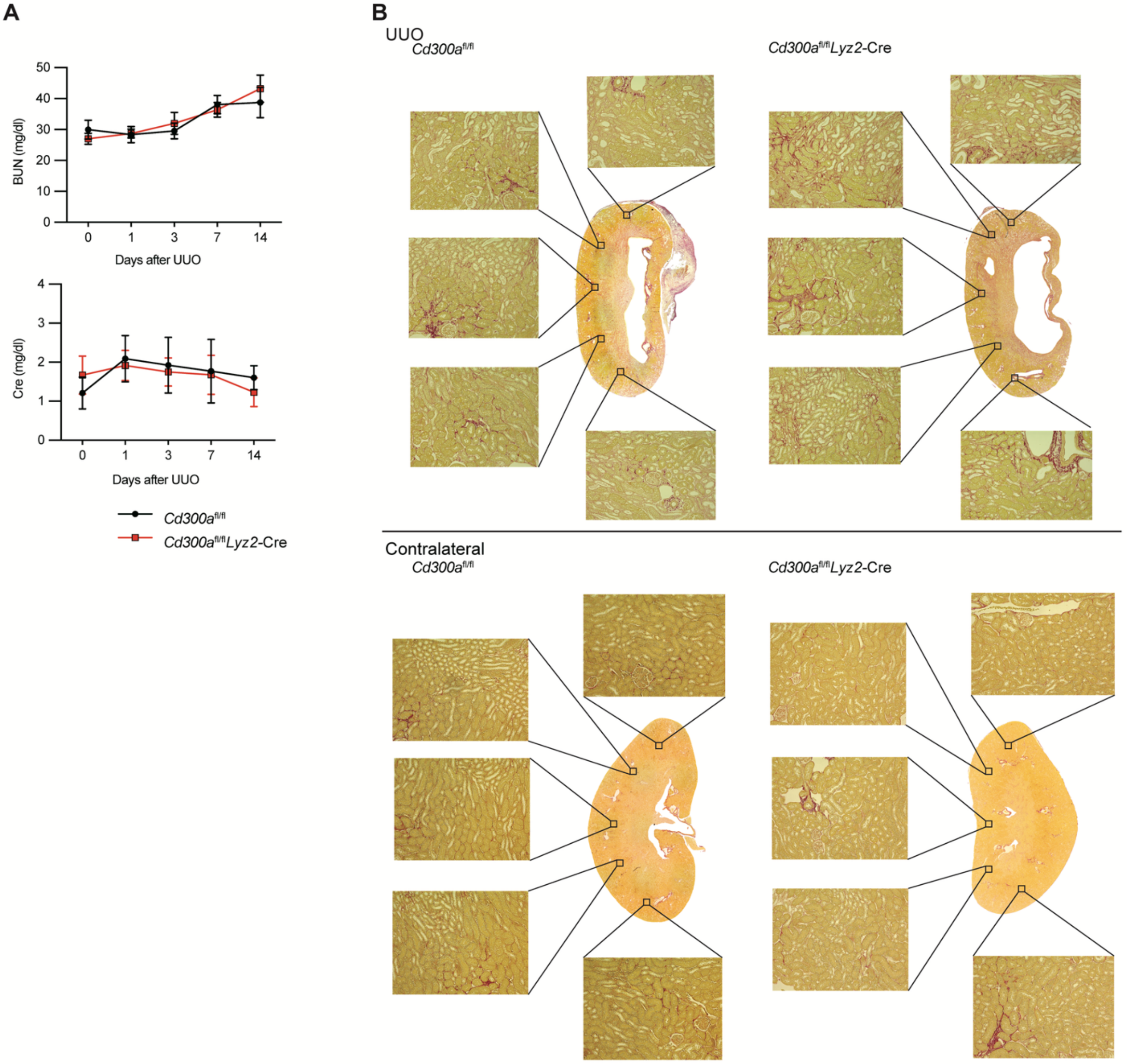


**Supplementary Fig. 1**

(A) Concentration of plasma blood urea nitrogen (BUN) and creatinine (Cre) were analyzed before (0 d) and 1, 3, 7 and 14 d after UUO in *Cd300a*$^{fl/fl}$ (n=12 [d 0], 8 [d 1],8 [d 3], 19 [d 7], 11 [d 14]) and *Cd300a*$^{fl/fl}$*Lyz2*-Cre (n=14 [d 0], 7 [d 1],7 [d 3], 11 [d 7], 4 [d 14]) mice. (B) Representative Sirius-red staining of whole kidneys on 14 d after UUO. Five regions were selected from the entire kidney for analysis, and the final quantification is presented as their average value.

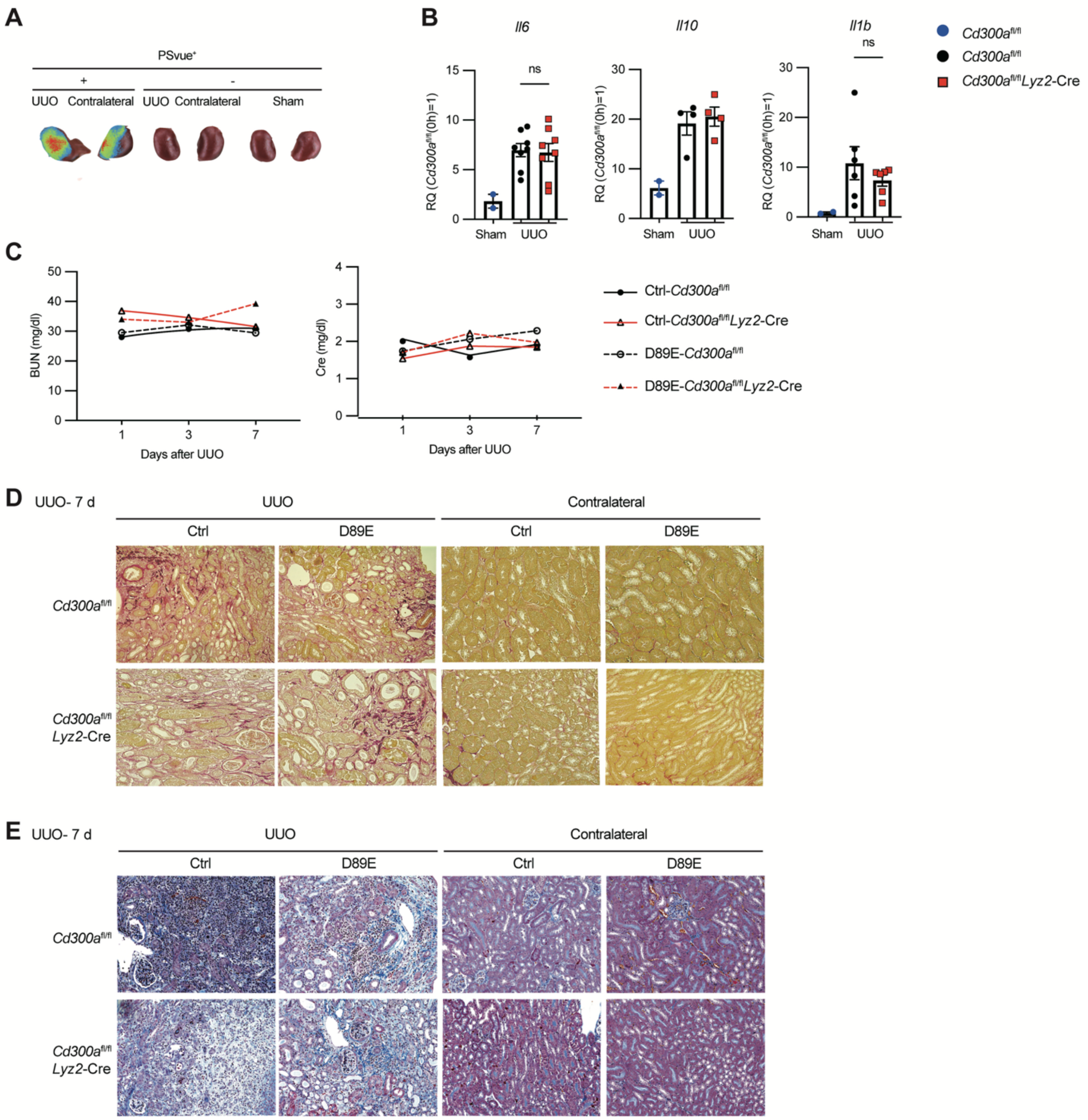


### Supplementary Fig. 2

(A) Flag-tagged D89E-MFGE8 was administered intravenously and just after UUO and kidney section was analyzed staining by anti-Flag antibody to detect the dead cells. (B) Q-PCR analysis of *Il6, Il10* and *Il1b* expressions in the kidneys 3 d after UUO in *Cd300a*$^{fl/fl}$ and *Cd300a*$^{fl/fl}$*Lyz2*-Cre mice. (C) Plasma concentrations of BUN and Cre 1, 3 and 7 d after UUO in EPT-D89E-MFGE8 (Ctrl) or D89E-MFGE8 (D89E) administered to *Cd300a*$^{fl/fl}$ (n=7 [d 1],4 [d 3], 3 [d 7] in control group; n=7 [d 1],4 [d 3], 4 [d 7] in D89E group) and *Cd300a*$^{fl/fl}$*Lyz2-Cre* (n=6 [d 1],3 [d 3], 3 [d 7] in control group; n=5 [day 1],3 [day 3], 3 [d 7] in D89E group) mice after UUO. (D and E) Representative Sirius-red (D) or Masson- trichrome (E) staining in kidneys from *Cd300a*$^{fl/fl}$ and *Cd300a*$^{fl/fl}$*Lyz2*-Cre mice administered with Ctrl or D89E at 7 d after UUO. Scale bars, 100 µm. Data are presented as mean ± SEM. Statistics analyses performed using one-way ANOVA (B). ns, not significant.

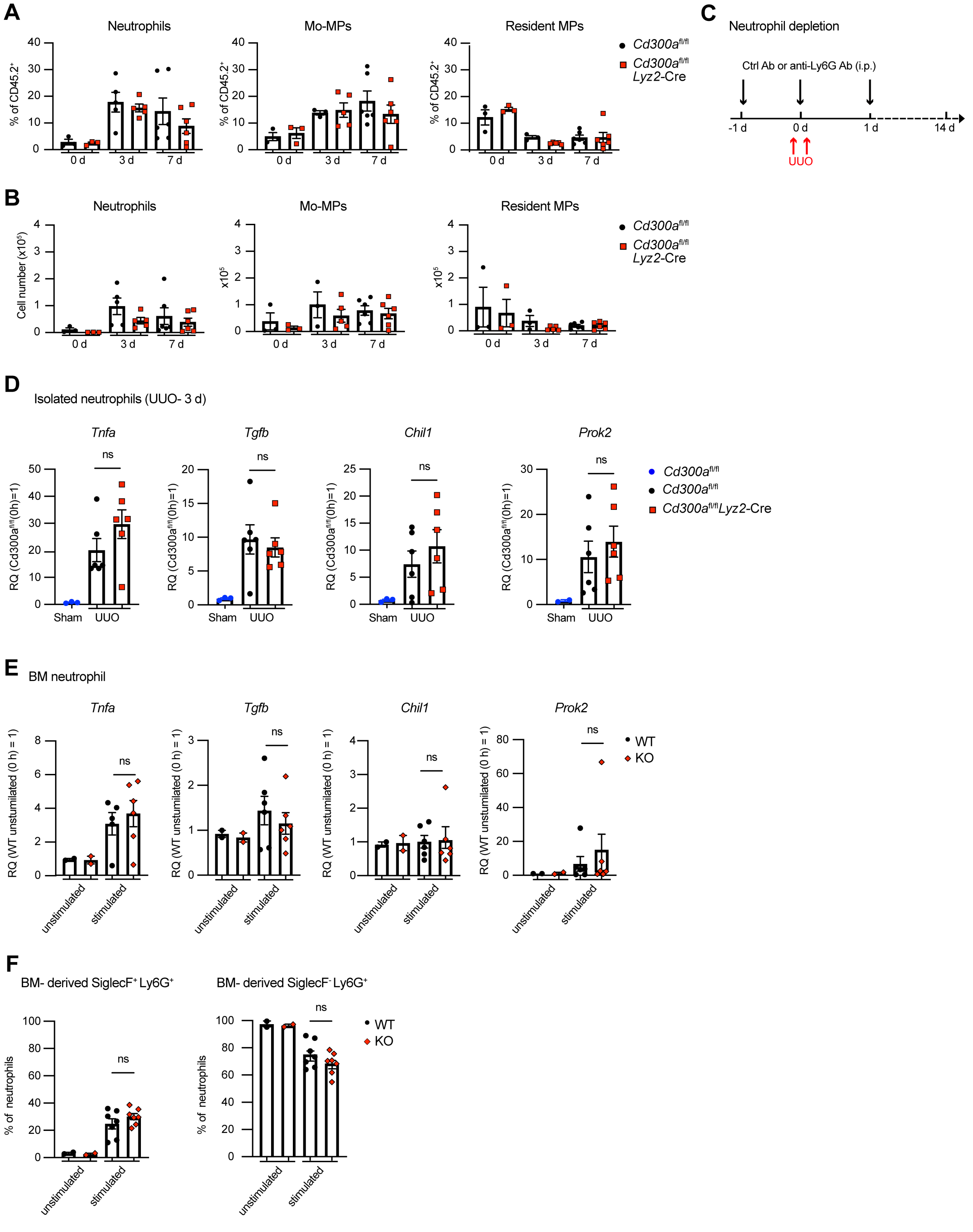


**Supplementary Fig. 3**

(A and B) Quantitative data of the percentage (A) and number (B) of neutrophils, monocyte-derived macrophages (Mo-MPs) and renal resident macrophages (resident MPs) 0, 3 and 7 d after UUO in

*Cd300a*$^{fl/fl}$ and *Cd300a*$^{fl/fl}$*Lyz2*-Cre mice. (C) Schematic diagram of neutrophil depletion by anti-Ly6G mAb. Control antibody (Ctrl Ab) or anti-Ly6G Ab was administered for three consecutive days, followed by UUO surgery on the next day. (D) Expression levels of *Tnfa*, *Tgfb*, *Chil1* and *Prok2* mRNA in isolated neutrophils from *Cd300a*$^{fl/fl}$ and *Cd300a*$^{fl/fl}$*Lyz2*-Cre mice at 3 d post UUO. (E) Bone marrow–derived (BM) neutrophils isolated from wild-type (WT) and *Cd300a*$^{-/-}$ mice were stimulated with injured kidney supernatants. Q-PCR analysis of *Tnfa*, *Tgfb*, *Chil1* and *Prok2* expressions. Data are presented as mean ± SEM. Statistics analyses performed using one-way ANOVA (A, B, D and E). ns, not significant.

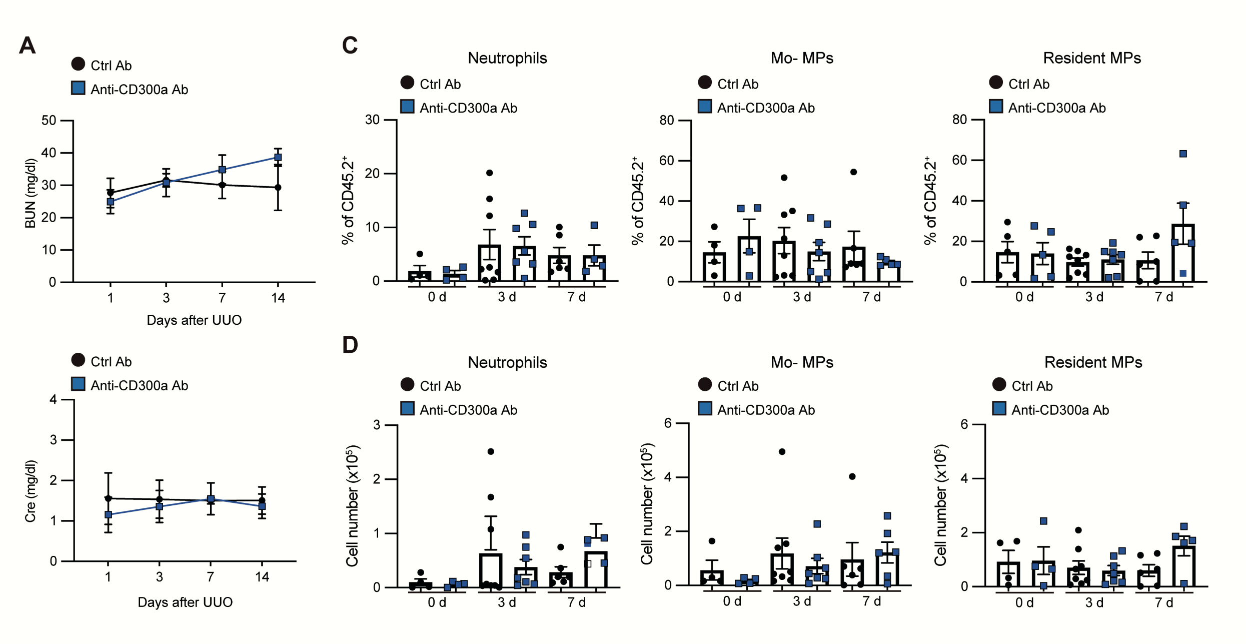


**Supplementary Fig. 4**

WT mice were treated with control antibody (Ctrl) or anti-CD300a antibody (Anti-CD300a) 2 h before UUO. (A) Concentration of BUN and Cre were analyzed 1, 3, 7 and 14 d after UUO. (B and C) Quantitative data of the percentage (B) and number (C) of neutrophils, Mo-MPs and resident MPs 0, 3 and 7 d after UUO in ctrl or anti-CD300a mAb treated mice.
